# Volitional cocaine taking engages distinct medium spiny neuron and astrocyte transcriptional programs in the rat nucleus accumbens

**DOI:** 10.64898/2026.06.19.733392

**Authors:** Heath D. Schmidt, Richard C. Crist, Samar N. Chehimi, Riley Merkel, Morgan Faist, Vaishnavi Joshi, Jackson E. Shuey, Benjamin C. Reiner

## Abstract

Cocaine use disorder (CUD) remains a major public health concern with no FDA-approved pharmacotherapy, underscoring the need to define the cellular and molecular adaptations produced by voluntary cocaine taking. The nucleus accumbens (NAc) is a key substrate for cocaine reinforcement and drug-seeking behavior, but interpretation of the functional role of its cellular heterogeneity in these behaviors is limited by past bulk transcriptomic studies. Here, we used single-nucleus RNA sequencing to profile the NAc of male and female rats that self-administered intravenous cocaine for 10 consecutive days versus yoked saline controls. After quality control, we analyzed 36,766 nuclei spanning major neuronal, glial, and vascular cell populations. Pseudobulk differential-expression analyses identified 478 cocaine-associated cell type-specific transcriptional changes that were concentrated in discrete medium spiny neuron (MSN) subclasses and astrocytes. D1 *Ebf1*+ MSNs showed the largest transcriptomic response, accounting for ∼40% of all differential-expression events, followed by D2 *Stk32a*+ MSNs, astrocytes, and D1 *Ppm1e*+ MSNs. These responses were largely cell type-specific, indicating that cocaine self-administration engages multiple molecular programs rather than a uniform accumbens-wide transcriptional signature. Immediate-early gene module-score analyses further revealed cocaine-associated activation states in select neuronal and non-neuronal cell populations, including D1 *Ebf1*+ MSNs, Drd3+ neurons, Sst+ interneurons, astrocytes, and oligodendrocytes. Gene-set, pathway, and upstream-regulator analyses nominated synaptic organization, axon guidance, RAS/MAPK signaling, NMDA receptor-associated signaling, and CREB-related transcriptional regulation as candidate mechanisms of cocaine-evoked plasticity. Together, these data provide a cell type-resolved resource for understanding how voluntary cocaine taking alters the rat NAc transcriptome and identifies discrete neuronal and glial cell populations for future mechanistic studies using preclinical CUD models.

## Introduction

Cocaine use disorder (CUD) remains an urgent and mechanistically unresolved public health concern. In 2024, 4.3 million people in the United States aged 12 years or older reported past-year cocaine use. While the prevalence of cocaine use amongst adults 26 years of age or older has remained constant, the percentage of adolescents (12 to 17 years of age) using cocaine significantly increased from 2021 to 2024 [1]. The incidence of fatal cocaine overdose increased substantially over the preceding decades and remains a major contributor to stimulant-related mortality [2, 3]. Yet, despite decades of clinical and preclinical research, there are currently no FDA-approved pharmacotherapies to treat CUD [4]. This therapeutic gap reflects, in part, an incomplete understanding of the molecular adaptations produced by voluntary cocaine taking in the neural circuits that regulate reinforcement, cue learning, motivation, and relapse.

The nucleus accumbens (NAc) is a central node in neural circuits regulating the rewarding and reinforcing effects of addictive drugs. Cocaine elevates dopaminergic signaling in mesolimbic and striatal neural circuitry, and the NAc integrates dopaminergic signals with cortical, limbic, and thalamic glutamatergic inputs to shape motivated behavior and drug seeking [5, 6]. The principal output neurons of the NAc are GABAergic medium spiny neurons (MSNs), which were classically categorized into dopamine receptor D1R- and D2R-expressing populations; D1 and D2 MSNs, respectively. That framework proved useful for examining D1 and D2 MSN manipulations, which produced distinct, and in some contexts, opposing effects on cocaine reward and cocaine-induced plasticity [7, 8]. However, evidence suggests the D1/D2 dicotomy provides an incomplete understanding of the role of NAc MSNs during hedonic behaviors. A recent single-nucleus atlas of the rat NAc resolved multiple molecularly distinct MSN subclasses beyond broad dopamine receptor identity, with some corresponding to the matrix and striosomal comparts of the NAc [9]. While some studies suggest that D1R- and D2R-expressing MSNs have opposing roles in cocaine-mediated behaviors, this picture is likely more complex; varying by stimulation pattern, extinction and withdrawal state, projection target, and molecular subtype [10-14]. Furthermore, cocaine-induced adaptations are not restricted to MSNs. Astrocytes and other glial populations regulate extracellular glutamate, synaptic homeostasis, and drug-seeking behavior in NAc circuits [15, 16]. Thus, understanding CUD-relevant plasticity requires moving beyond major cell types to identify the specific neuronal and non-neuronal cell populations engaged by cocaine experience.

Early transcriptomic studies using microarrays and bulk RNA sequencing established that cocaine alters gene expression across rodent, non-human primate, and human brains [17, 18]. While these studies were foundational to our understanding of the effects of cocaine on the neural transcriptome, the bulk tissue approach averages expression across a heterogeneous mixture of molecularly distinct glial, vascular, and neuronal cell populations. Consequently, bulk transcriptomics can identify broadly altered molecular mechanisms, but fails to assign cell type specificity and are unable to identify more subtle alterations in transcriptomic profiles. Single-cell and single-nucleus RNA sequencing (snRNA-seq) overcomes these limitations by simultaneously defining molecular-distinct cell populations and quantifying cell type-specific transcriptional responses in heterogeneous brain tissue [19]. This resolution is especially important for addiction neuroscience, where molecular changes in rare or functionally- and connectively-specialized cell populations may be obscured in bulk transcriptome profiles. Recent human CUD multi-omics studies further emphasize the value of this approach, implicating cell type-specific striatal alterations in MSNs and astrocytes [20, 21].

In rats, acute cocaine induces cell type-specific immediate-early gene programs in striatal neurons [22], while acute and repeated experimenter-administered cocaine identified a single D1 MSN subpopulation that was transcriptionally responsive [23]. In mice, fluorescence-activated nuclear sorting coupled with snRNA-seq also revealed distinct D1 and D2 transcriptome alterations associated with acute cocaine exposure [11]. These studies demonstrate the power of cell type-resolved approaches, but most have relied on non-contingent drug exposure or sorted neuronal populations. Intravenous cocaine self-administration is a translationally relevant model of volitional drug taking that may recruit molecular programs that are not fully captured by passive cocaine exposure. Thus, there is a need for snRNA-seq studies of the NAc from animals self-administering cocaine to identify cell type-specific changes in gene expression associated with volitional drug-taking behavior.

Here, we used snRNA-seq to profile the NAc transcriptomes of male and female rats that self-administered intravenous cocaine versus their saline yoked controls. We identified all major neuronal and non-neuronal cell populations in the rat NAc and mapped cocaine-associated differential expression across these populations. The strongest transcriptional responses were observed in discrete MSN subclasses as well as astrocytes. Enrichment and upstream-regulator analyses nominated synaptic, axon-guidance, circadian, RAS/MAPK, NMDA receptor, and CREB-related programs as candidate mechanisms of cocaine-evoked plasticity. Together, these data provide a cell type-resolved resource for understanding how voluntary cocaine taking reshapes the rat NAc transcriptome and identifies specific neuronal and glial cell populations for follow-up mechanistic studies.

## Materials and Methods

### Drugs

Cocaine HCl was obtained from the National Institute on Drug Abuse (Rockville, MD) and dissolved in bacteriostatic 0.9% saline. The dose of cocaine self-administered was based on previously published studies in rats from our group [24-27].

### Animals and housing

Male and female Sprague-Dawley rats (Rattus norvegicus) weighing 225-250g were obtained from Taconic Laboratories (Rensselaer, NY). Rats were housed individually on a 12/12 h light/dark cycle and maintained on ad libitum food and water. All behavioral procedures were performed during the light cycle. All experimental protocols were consistent with the guidelines issued by the U.S. National Institutes of Health and were approved by the Institutional Animal Care and Use Committee (IACUC) of the University of Pennsylvania.

### Catheterization Surgery

Rats were handled daily and allowed one week to acclimate to their home cages upon arrival. Prior to surgery, rats were anesthetized with 100 mg/kg ketamine and 10 mg/kg xylazine (Covetrus). An indwelling catheter (SAI Infusion Technologies, Lake Villa, IL) was inserted into the right jugular vein and sutured in place. The catheter was routed subcutaneously to a mesh backmount that was implanted dorsal to the shoulder blades in the mid-scapular region. To prevent infection and maintain patency, catheters were flushed daily with 0.2 ml of the antibiotic Timentin (0.93 mg/ml; Fisher, Pittsburgh, PA) dissolved in heparinized 0.9% saline (Butler Schein, Dublin, OH). When not in use, catheters were sealed with plastic obturators.

### Cocaine self-administration

Following seven days of recovery, rats were randomly assigned to one of two treatment groups: cocaine-experimental or yoked saline controls. Cocaine-experimental rats were placed into operant conditioning chambers and allowed to lever-press for intravenous infusions of cocaine (0.25 mg/kg/infusion, infused over 5-s) on a fixed-ratio 1 (FR1) schedule of reinforcement, similar to our previous studies of cocaine self-administration in rats [24-27]. All self-administration sessions were two hours in duration and were conducted over 10 consecutive days. Each cocaine infusion was paired with a 20-s contingent light cue illuminated directly above the active lever (i.e., drug-paired lever). A 20-s time-out period followed each cocaine infusion, during which time active lever responses were tabulated but had no scheduled consequences. Response made on the inactive lever, which had no scheduled consequences, were also recorded during the self-administration sessions. Each cocaine-experimental rat was paired with a yoked rat that received infusions of saline. While lever pressing for the yoked saline rats had no scheduled consequences, these rats received the same number and temporal pattern of infusions as self-administered by their paired cocaine-experimental rat. Rats were euthanized immediately after their last self-administration session. Whole brains were dissected, flash frozen in -20°C isopentane and stored at -80°C.

### Nuclei Isolation, Library Preparation, and Sequencing

Bilateral NAc tissue was punched from frozen brains and nuclei suspensions were prepared, using our published methods [28-32]. The nuclei suspensions were used as input to the 10x Genomics Multiome assay, per manufacturer protocols. Resulting libraries were sequenced on an Illumina NovaSeq 6000, and reads were deconvoluted and aligned to rat genome (rn7.2.110) using Cell Ranger ARC (v2.0.2).

### Quality Control and Clustering of snRNA-seq Data

Seurat v5.2.1 was used to perform quality control and clustering. Due to a lack of nuclei with sufficient depth of ATAC reads, the ATAC data was dropped from subsequent analyses, and the data were analyzed as snRNA-seq only. Nuclei with ≤200 captured genes or ≥5% mitochondrial transcript reads were removed. SoupX was then used to correct for ambient RNA contamination. Nuclei with abnormally low (≤200) or high (≥5000) numbers of captured genes after ambient RNA correction were removed. The count data were normalized and scaled with SCTransform, which also regressed out the percentage of mitochondrial transcripts. Samples were then integrated to ensure consistent cluster identification across samples and experimental groups. Doublets were identified with scDblFinder v1.16.0. Clusters with >50% doublets and all individual nuclei identified as doublets were removed. Finally, any remaining cell populations with low numbers of unique molecular identifiers (UMIs) or mixed markers for major cell types were removed. The remaining clusters in the cleaned dataset were assigned to major cell types based on expression of curated known marker genes [9].

### Differential expression and gene set and pathway analysis

Pseudobulk differential expression between cases and controls was performed in each cell population separately using EdgeR LRT. Sex was used a covariate in the primary combined analysis. Males and females were also analyzed independently. Cell type specific differential expressed gene list were used as inputs FUMA and IPA as we previously described [28, 29, 33, 34]. Additional gene set analysis and network diagramming was performed with Metascape [35]. Immediate early gene module scores were calculated using the AddModuleScore function in Seurat, using 25 immediate early genes (*Arc, Bdnf, Cdkn1a, Dnajb5, Egr1, Egr2, Egr4, Fos, Fosb, Fosl2, Homer1, Junb, Nefm, Npas4, Nr4a1, Nr4a2, Nr4a3, Nrn1, Ntrk2, Rheb, Sgsm1, Syt4*, and *Vgf*). A control feature score of 5 was used, and differences in module scores between the cocaine and saline groups waswere assessed using a Wilcoxon Rank Sum test in SCpubr [36].

### Data and code availability

The single nuclei sequencing and sample metadata are available at the NCBI Gene Expression Omnibus (GEO) under accession number ########. The Seurat object and all code are available from the corresponding author by request.

## Results

### Cocaine self-administration produces robust operant responding in male and female rats

Male and female Sprague-Dawley rats (n = 16, 4 rats/sex/condition) were surgically implanted with indwelling jugular catheters and, after recovery, were randomized to cocaine self-administration or yoked saline control groups (**Figure 1A**). Rats acquired cocaine self-administration and escalated intake over 10 days, as shown in **Figures 1B-D**. Micropunches containing the NAc core and shell were taken for nuclei isolation (**Figure 1E**).

**Figure 1:**
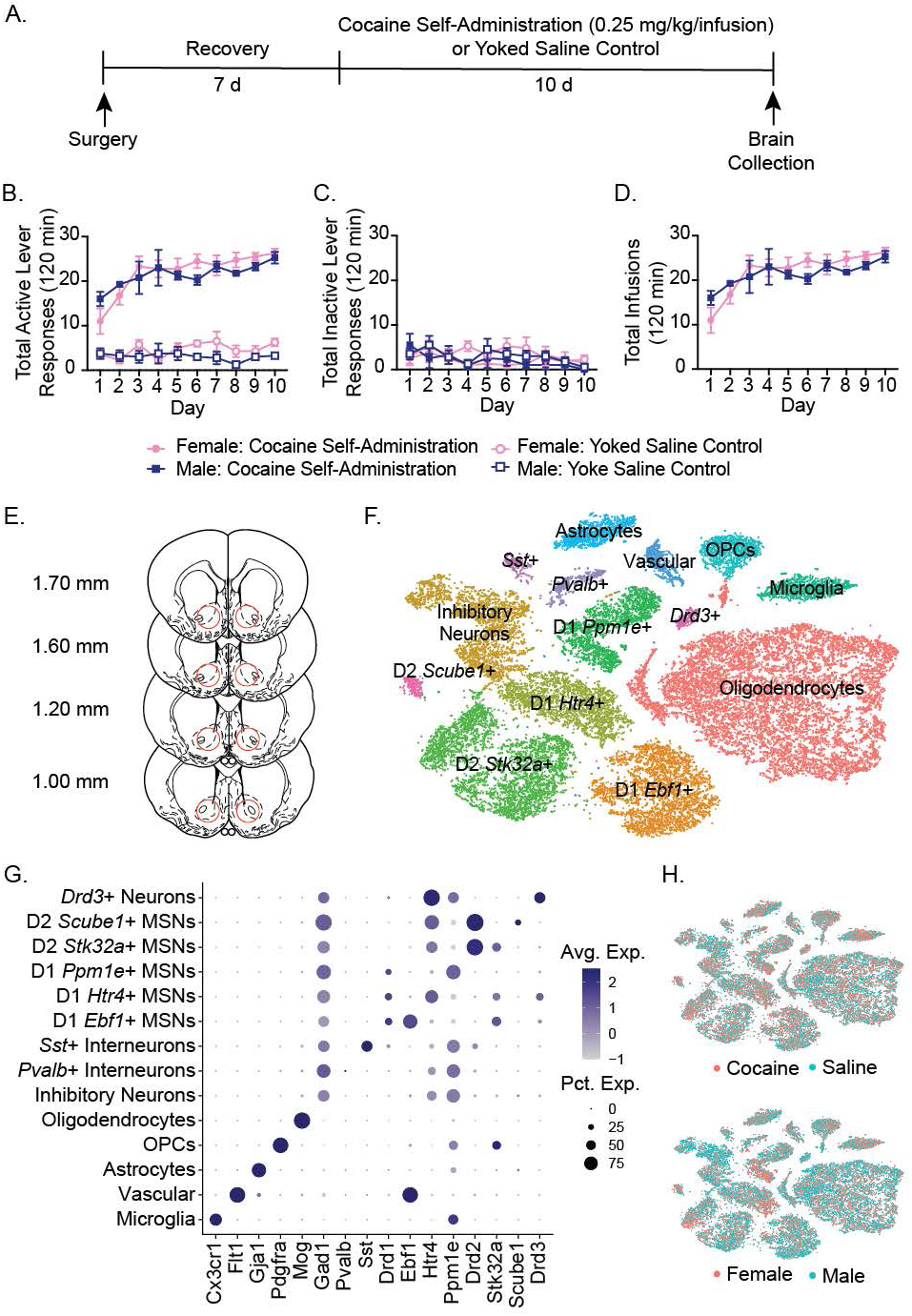
Clustering of snRNA-seq data from cocaine self-administration paradigm. Male and female Sprague-Dawley rats were randomized to cocaine self-administration or yoked saline control groups (n = 4/sex/group). (a) Timeline of jugular catheter surgery, recovery, and self-administration protocol. (b-d) Line graphs of self-administration data demonstrate that male and female rats in the cocaine group escalate active lever presses and infusions over 10 days. (e) Red circles indicate the location of the tissues punches taken from the rat NAc. (f) T-distributed stochastic neighbor embedding (t-SNE) dimension reduction plot with nuclei colored by major cell type. (g) Dot plot showing expression of marker genes for known cell types in the NAc. (h) T-SNE plots colored by group or sex demonstrate successful integration of all samples.

### Cocaine self-administration induces cell type-specific transcriptional remodeling concentrated in discrete MSN subtypes and astrocytes

After quality control, ambient RNA correction, doublet removal, and filtering of low-quality populations, the final dataset contained 36,766 NAc nuclei. Unsupervised clustering followed by t-distributed stochastic neighbor embedding showed that nuclei were separated into the major neuronal, glial, and vascular cell populations expected in the rat NAc, including molecular defined subclasses of MSNs (**Figure 1F**). Cluster annotations were further confirmed by canonical marker-gene expression (**Figure 1G**). The integrated dataset showed broad representation of both treatment groups and both sexes across the clusters, with no obvious segregation of nuclei by cocaine exposure or sex in the low-dimensional embedding (**Figure 1H**). This integration supported subsequent cell type-resolved comparisons between cocaine-experienced rats and yoked saline controls.

We next performed pseudobulk differential-expression analysis to identify cocaine-associated transcriptional changes within each annotated cell population. The primary analysis of combined males and female, using sex as a covariate, identified 478 cell type-specific differential-expression events between cocaine-experienced rats and yoked saline controls. These changes were not uniformly distributed across the NAc cellular landscape. Instead, the majority of differential expression was concentrated in four cell populations: D1 *Ebf1*+ MSNs, D2 *Stk32a*+ MSNs, astrocytes, and D1 *Ppm1e*+ MSNs (**Figure 2A–D**; **Table S1**). D1 *Ebf1*+ MSNs showed the largest cocaine-associated transcriptional response, with 192 differentially expressed genes (DEGs), accounting for approximately 40% of all differential-expression events detected in the combined analysis (**Figure 2A**). Within this MSN cell population, upregulated genes in the context of cocaine self-administration included *Hectd2, Irs2, Bcl11b, Traf3, Per2*, and *Ubash3b*, whereas downregulated genes included *Itgav, Cntn1, Samd5*, and *Pacs1*. Thus, the largest cocaine-responsive MSN cell population contained changes in genes related to transcriptional regulation, intracellular signaling, adhesion, and synaptic/cellular architecture.

**Figure 2:**
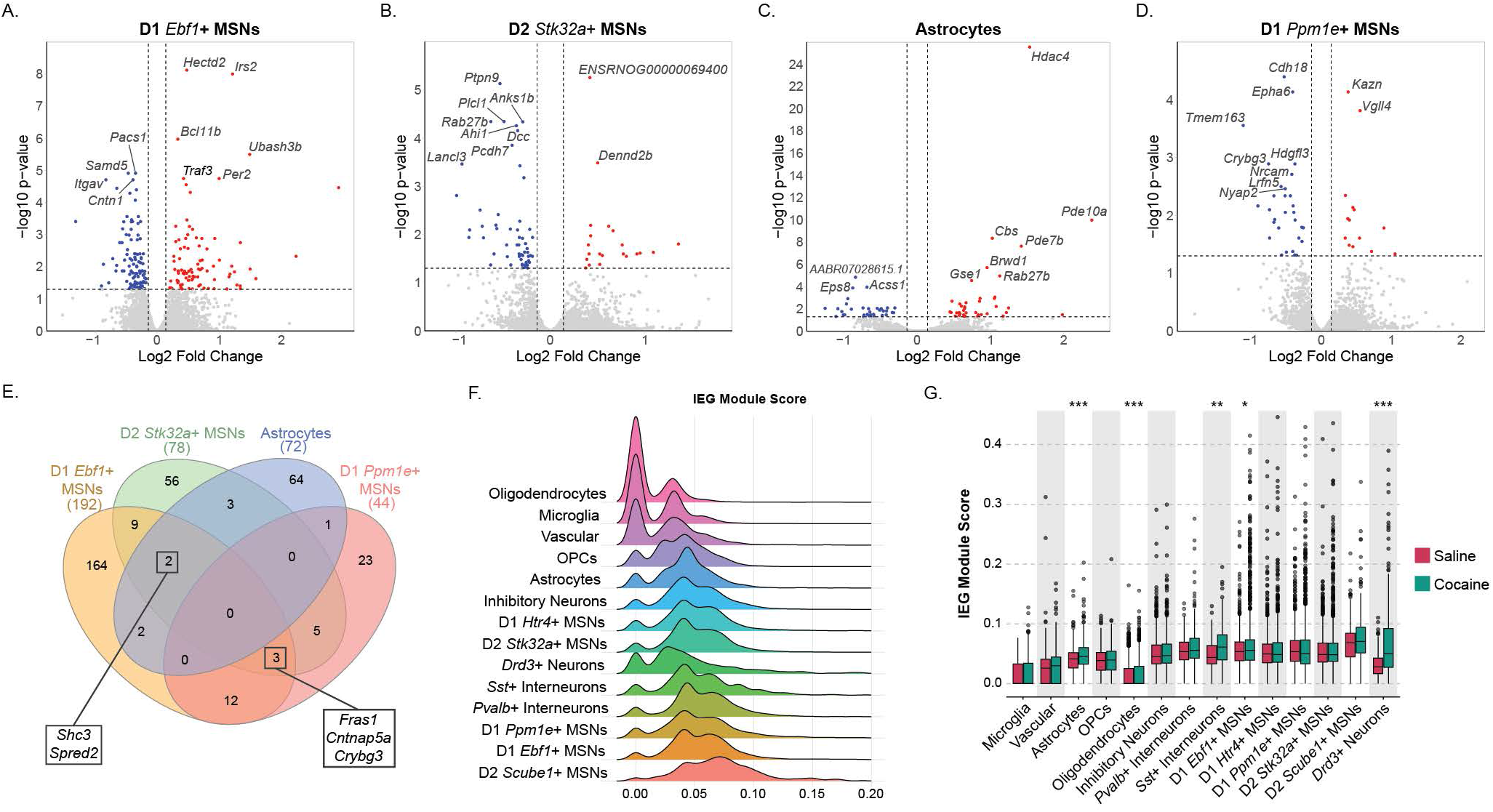
Cell type-specific differential expression following cocaine self-administration. (a-d) Volcano plots visualize the results of pseudobulk differential expression analysis with EdgeR for the four cell types with the most DEGs. The ten genes with the smallest p-values are labeled on each plot. (e) A Venn diagram indicates the number of shared DEGs between these four cell types. Genes differentially expressed in three cell types are highlighted. (f) Ridge plot showing IEG expression for nuclei in all clusters, represented by a module score derived from expression of 25 IEGs. (g) Box and whisker plots indicating IEG module scores for the cocaine and saline groups in each cell type. Dots represent nuclei with module scores more than 1.5x the inter-quartile range from the edges of the boxes. Asterisks denote results of Wilcoxon Rank Sum tests comparing module scores between the two groups. *, p < 0.05; **, p < 0.01; ***, p < 0.005

D2 *Stk32a*+ MSNs contained the second-largest cocaine-associated transcriptional response, with 78 DEGs (**Figure 2B**). In this cluster, the strongest downregulated genes included *Ptpr9, Anks1b, Rab27b, Ahi1, Dcc, Pcdh7*, and *Lancl3*, whereas upregulated genes included *Dennd2b* and an unannotated transcript, ENSRNOG00000069400. Astrocytes contained 72 DEGs and showed several of the most statistically significant individual cocaine-associated genes in the dataset (**Figure 2C**). *Hdac4* was the most significant DEG identified in astrocytes, with additional upregulated genes including *Pde10a, Pde7b, Cbs, Brwd1, Gse1*, and *Rab27b*. Downregulated genes in astrocytes included *Eps8, Acss1*, and *AABR07028615*.*1*. D1 *Ppm1e*+ MSNs contained 44 DEGs, including upregulation of *Kazn* and *Vgll4* and downregulation of *Cdh18, Epha6, Tmem163, Crybg3, Hdgfl3, Nrcam*, and related adhesion-associated genes (**Figure 2D**). DEGs amongst the four most cocaine-responsive cell populations (i.e., D1 *Ebf1*+ MSNs, D2 *Stk32a*+ MSNs, Astrocytes, and D1 *Ppm1e*+ MSNs) were largely transcriptomically distinct with only select convergent transcriptional events across MSN subtypes and astrocytes (**Figure 2E**).

We next performed an exploratory sex-stratified differential-expression analysis since both males and females were included in the primary analysis. The male-only analysis (n = 4/group) identified 520 cell type-specific differential-expression events, whereas the female-only analysis (n = 4/group) identified 55 events (**Tables S2 and S3**). The greater number of DEGs detected in males should be interpreted cautiously because female samples contributed fewer nuclei to the final dataset than male samples (13,292 vs 23,474 nuclei, respectively), reducing power to detect female-specific effects. Nevertheless, the sex-stratified analyses recovered a subset of events identified in the primary combined model: 168 male-only events and 27 female-only events overlapped with the combined analysis, and nine events were detected in both sex-stratified analyses.

### Cocaine self-administration increases immediate-early gene module scores in select neuronal and glial populations

To complement the gene-level differential-expression analyses, we next quantified cellular activation states using polygenic modeling of cell activation for a composite score of 25 immediate early genes. Across all nuclei, IEG module-score distributions differed substantially by cell population (**Figure 2F**). D2 *Scube1*+ MSNs, D1 *Ebf1*+ MSNs, and D1 *Ppm1e*+ MSNs showed the highest median IEG module scores, indicating that these MSN subtypes express relatively high levels of activity-regulated transcripts at the time of tissue collection, independent of treatment group. Direct comparison of cocaine-experienced rats and yoked saline controls within each cell population revealed significant cocaine-associated increases in IEG module scores in both neuronal and non-neuronal populations (**Figure 2G**). Among glial populations, astrocytes and oligodendrocytes showed significantly higher IEG module scores after cocaine self-administration. Among neuronal populations, Sst+ interneurons, D1 *Ebf1*+ MSNs, and Drd3+ neurons showed significant cocaine-associated increases. Notably, the populations with significant IEG module-score changes only partially overlapped with the populations carrying the largest DEG burdens. For example, D1 *Ebf1*+ MSNs and astrocytes showed both prominent differential expression and significant IEG score increases, whereas oligodendrocytes showed a significant IEG score increase despite not being among the top DEG-containing clusters. Conversely, D2 *Stk32a*+ MSNs and D1 *Ppm1e*+ MSNs showed substantial differential expression but did not exhibit comparably strong cocaine-associated IEG module-score shifts. Thus, cocaine-associated transcriptional remodeling and immediate-early gene activation identify overlapping but non-identical dimensions of cellular response in the NAc.

### D1 Ebf1+ MSN DEGs are enriched for functional gene sets and upstream intracellular signaling pathways

Since D1 *Ebf1*+ MSNs contained the largest number of cocaine-associated DEGs, we first examined functional enrichment within this cell population. Gene-set enrichment analysis revealed a broad, interconnected network of biological processes (**Figure 3A**; **Table S4**). These enrichments indicate that cocaine-associated DEGs in D1 *Ebf1*+ MSNs converge on molecular programs central to synaptic organization, intracellular signal transduction, cell adhesion, and striatal neuronal identity. We next tested whether DEGs from the four most affected cell populations were enriched for known transcription factor and microRNA target sets (**Table S5**). Transcription factor target enrichment was strongest and most diverse in D1 *Ebf1*+ MSNs (**Figure 3B**). Astrocytes and D2 *Stk32a*+ MSNs showed more restricted enrichment profiles, whereas D1 *Ppm1e*+ MSNs showed comparatively modest transcription factor target enrichment. MicroRNA target enrichment showed a similarly cell type-specific structure (**Figure 3C**). D1 *Ebf1*+ MSN DEGs were more robustly enriched for microRNA targets, compared to the other transcriptionally active cell types. Taken together, these results suggest that the most cocaine-responsive cell types are not only distinguished by DEG number but also by distinct upstream regulatory architectures.

**Figure 3:**
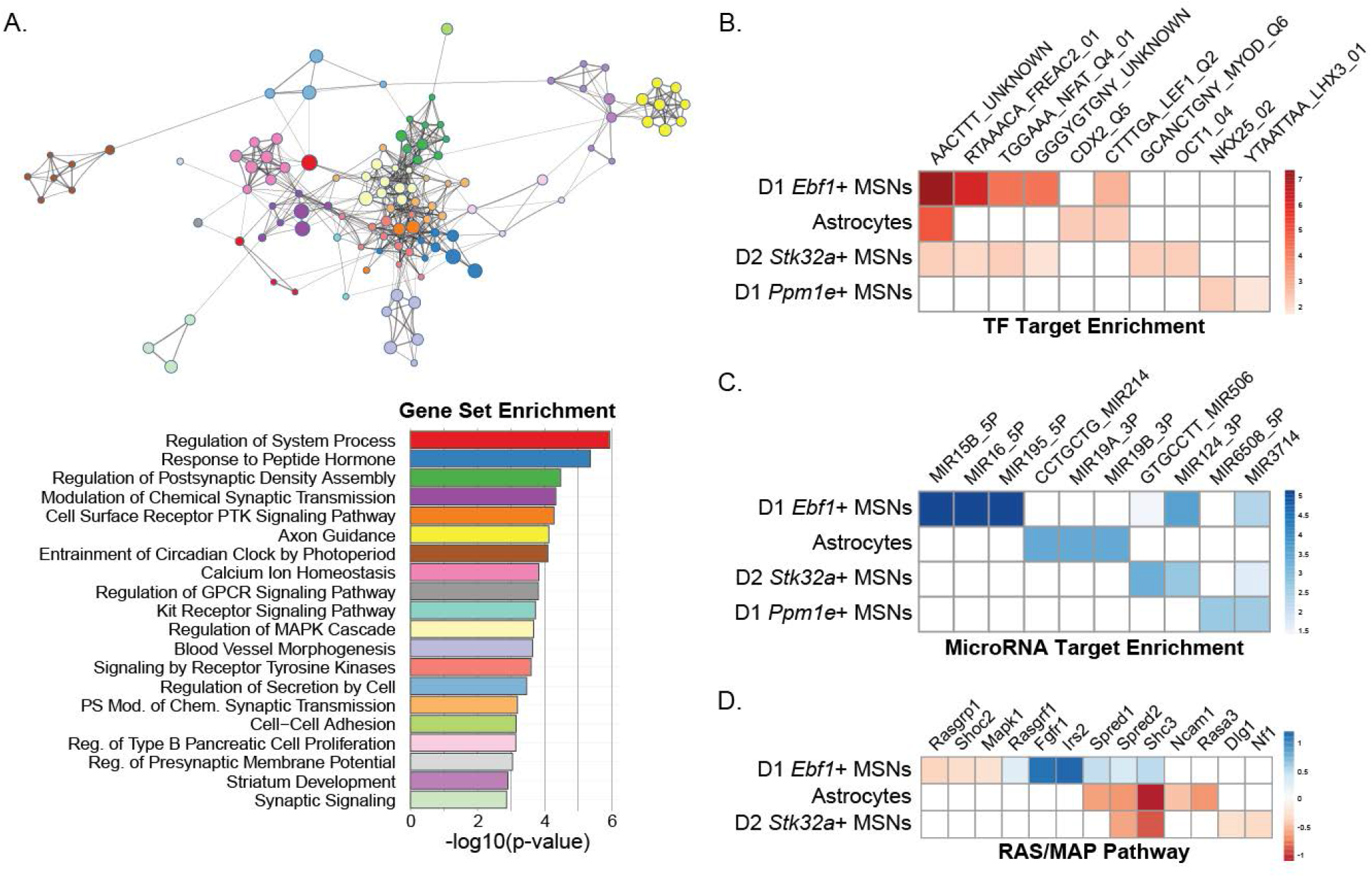
Enrichment of gene sets, regulatory targets, and biological pathways following cocaine self-administration. (a) Cytoscape network plot of enriched gene sets in DEGS from D1 *Ebf1*+ MSNs. Node size indicates -log10(p-value) of the hypergeometric test in Metascape after FDR correction. Shared node colors indicate closely related gene sets and edge widths indicate the number of shared genes between sets. Bar plot shows the -log10(p-value) of a representative gene set for each color group in the network plot. FUMA was used to identify enriched transcription factor target motifs (b) and microRNA target genes (c) in DEGs from the four cell types with the most differential expression. Heatmaps indicate -log10(p-value) of the hypergeometric test. (d) Heatmap shows the logFC of DEGs in the RAS/MAP pathway from three clusters. IPA predicts the pathway to be upregulated in D1 *Ebf1*+ MSNs and downregulated in astrocytes and D2 *Stk32a*+ MSNs.

### Pathway and regulatory analyses reveal shared but directionally distinct cocaine-associated signaling

Canonical pathway analysis identified RAS/MAPK signaling as a notable convergent pathway across three of the four most affected cell populations (**Figure 3D**). Although the pathway was implicated in multiple cell types, the predicted direction of pathway activity differed by cell population, with activity consistent with increased signaling In D1 *Ebf1*+ MSNs and decreased signaling in astrocytes and D2 *Stk32a*+ MSNs. Several of the genes associated with the signaling alterations were also part of the limited set of shared DEGs amongst the active cell types. These findings suggest that cocaine self-administration engages RAS/MAPK-associated signaling across multiple NAc cell types, but that the transcriptional direction and inferred functional consequence of this pathway engagement is cell type-dependent. Upstream regulatory network analysis identified higher-order regulatory structures associated with cocaine-induced transcriptional remodeling (**Table S6**). One prominent example was a dense regulatory network in D1 *Ebf1*+ MSNs that organized around NMDA receptor signaling (Figure 4). The inferred regulatory architecture linked NMDA receptor signaling to intracellular cascades involving PI3K, ERK/MAPK, p38 MAPK, calcineurin, NFAT-family signaling, and CREB1. CREB1 was predicted to be activated and was connected to multiple observed DEGs, including activity-regulated transcription factors and signaling molecules. This network analysis therefore places the D1 *Ebf1*+ MSN cocaine-response program within a broader framework of glutamatergic receptor signaling, intracellular kinase/phosphatase cascades, and activity-dependent transcriptional regulation. Together with the differential-expression and enrichment analyses, these studies identify D1 *Ebf1*+ MSNs as a major cellular substrate of cocaine-evoked transcriptional plasticity in the rat NAc while also identifying astrocytes and additional MSN subtypes as important, molecularly distinct components of the ventral striatal response to voluntary cocaine taking.

**Figure 4:**
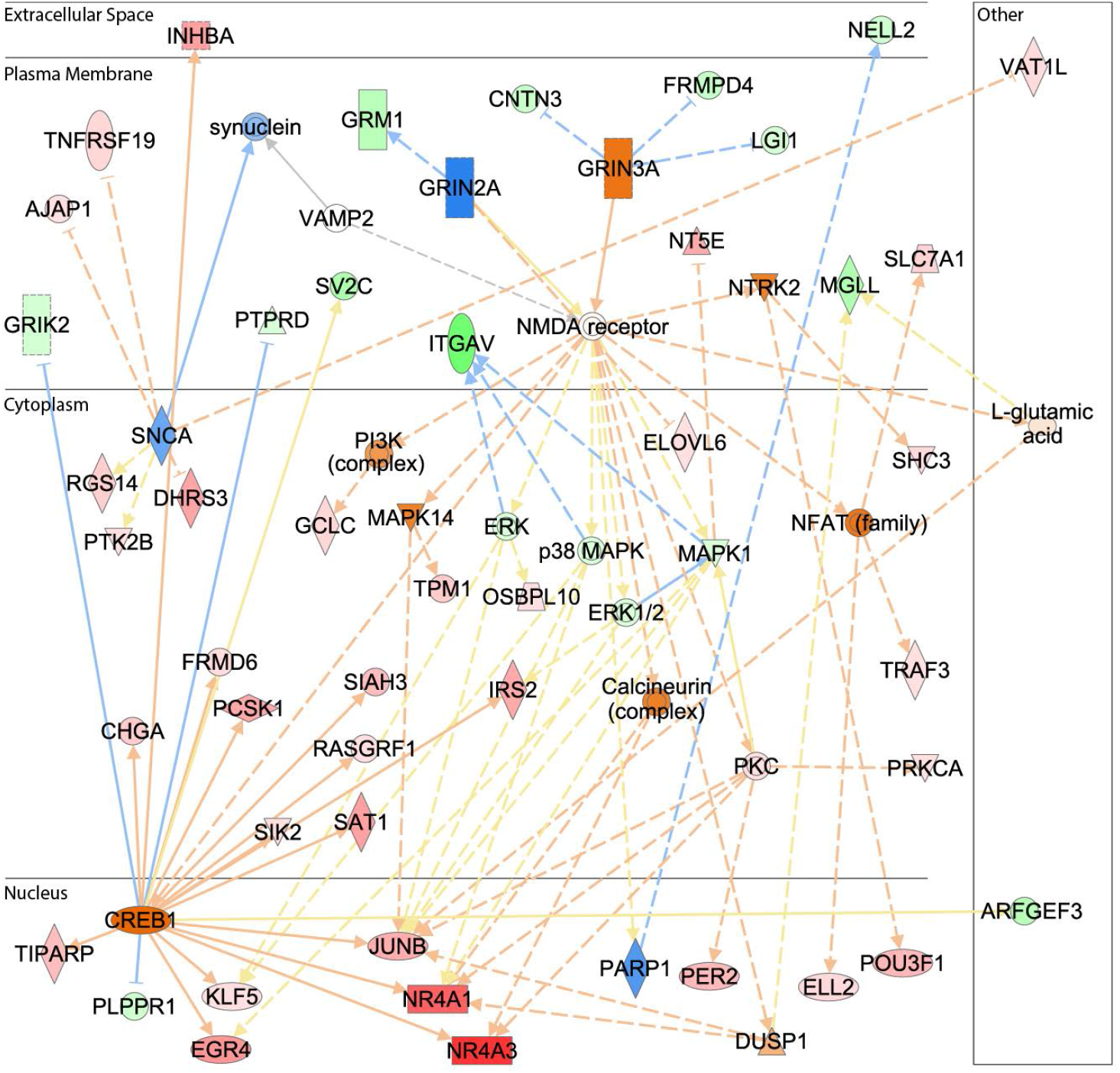
Dysregulation of a large regulatory network in D1 Ebf1+ MSNs following cocaine self-administration. Network diagram from IPA of observed differential expression and predicted functional changes based on DEGs from D1 *Ebf1*+ MSNs. Red and green molecules represent observed increased and decreased expression, respectively. Orange and blue molecules represent predicted increased and decreased function, respectively. Darker colors indicate larger predicted effect size

## Discussion

The present study expands our understanding of molecular basis of voluntary drug taking by identifying novel, cell type-specific transcriptomic changes in the NAc of cocaine experienced male and female rats. Specifically, we used single-nucleus RNA sequencing to resolve the major neuronal, glial, and vascular cell populations in the ventral striatum. We found that cocaine self-administration did not produce a uniform transcriptional response across the NAc. Instead, cocaine-associated DEGs were concentrated in a small number of molecularly defined cell populations, with the largest effects observed in D1 *Ebf1*+ MSNs, followed by D2 *Stk32a*+ MSNs, astrocytes, and D1 *Ppm1e*+ MSNs. The majority of DEGs in these cell populations were cell type-specific, indicating that voluntary cocaine taking engages multiple parallel transcriptional programs rather than a single pan-cellular cocaine signature. Immediate-early gene module-score analyses further showed that cocaine taking increased activity-associated transcriptional states in select neuronal and non-neuronal populations, including D1 *Ebf1*+ MSNs and astrocytes, while pathway analyses nominated synaptic organization, axon guidance, RAS/MAPK signaling, NMDA receptor-associated signaling, and CREB-related transcriptional regulation as candidate mechanisms of cocaine-evoked plasticity. Together, these findings identify discrete NAc cell populations and regulatory pathways that characterize the molecular foundation of voluntary cocaine taking.

A major strength of the present study is that it profiles the NAc after intravenous cocaine self-administration, a model of volitional drug taking. Prior single-nucleus studies showed that acute experimenter-delivered cocaine and repeated non-contingent cocaine exposure induced cell type-specific transcriptional responses in striatal neurons [11, 22, 23]. Those studies were essential for establishing that cocaine-responsive transcriptional programs can be resolved at single cell resolution. However, a significant limitation of these models is that non-contingent cocaine exposure and sorted neuronal populations cannot fully model the behavioral and motivational components of drug taking. The present dataset therefore fills an important gap by characterizing the cellular transcriptomic architecture of the NAc in male and female rats that self-administered cocaine. This complements recent work using single-nucleus multiomics in rats with divergent cocaine addiction-like behaviors, which identified amygdala adaptations after abstinence [37], as well as a human CUD multi-omics study showing cell type-specific striatal alterations in MSNs, astrocytes, and synaptic regulatory programs [20, 21]. The convergence of these studies supports the idea that substance use disorders are not simply associated with global transcriptional dysregulation, but with molecular remodeling in specific neuronal and glial cell populations embedded within distributed heterogenous addiction-relevant nuclei. Furthermore, these data suggest that future studies using an operant model to investigate transcriptomic changes during abstinence following volitional cocaine taking and examination of cocaine reinstatement-induced alterations of cell type-specific transcriptomes in the NAc are warranted.

The most striking cellular response in the current study was observed in D1 *Ebf1*+ MSNs. This cell population accounted for approximately 40% of all cocaine-associated differential-expression events in the combined analysis, showed increased IEG module scores after cocaine self-administration, and exhibited the strongest enrichment for transcription factor and microRNA target sets. These findings align with rat snRNA-seq studies showing that only a subset of Drd1-expressing MSNs expressing *Ebf1* is robustly transcriptionally responsive to non-contingent cocaine exposure [22, 23]. Importantly, the current results extend those observations to a volitional cocaine-taking paradigm, indicating that D1 *Ebf1*+ MSNs are a highly reproducible cocaine-responsive MSN subtype across experimental contexts, including a recent NAc MSN atlas effort [9]. The transcriptional programs altered in D1 *Ebf1*+ MSNs are especially compelling because they point toward molecular mechanisms that are central to cocaine-related synaptic plasticity, including processes related to structural and functional synaptic plasticity. Upstream-regulator analysis further identified a dense regulatory network organized around NMDA receptor signaling and connected to PI3K, ERK/MAPK, p38 MAPK, calcineurin, NFAT-family signaling, and CREB1. This is highly consistent with decades of work implicating glutamatergic signaling, intracellular kinase cascades, and activity-dependent transcription in cocaine-induced plasticity in the NAc [5, 6].

The effects of cocaine self-administration clearly extended beyond the D1 *Ebf1*+ MSN population, with D2 *Stk32a*+ MSNs and D1 *Ppm1e*+ MSNs also exhibiting substantial cocaine-associated differential expression, but with transcriptional profiles that were largely distinct from D1 *Ebf1*+ MSNs. This reinforces the growing view that NAc MSNs cannot be adequately described by a simple D1/direct versus D2/indirect pathway framework. NAc MSNs instead contribute to reward, aversion, cocaine seeking, and cocaine-induced plasticity in ways that depend on projection target, subregion, stimulation pattern, and molecular subtype [10-14]. In this context, the identification of cocaine-evoked transcriptional responses is MSN subtypes from both the matrix (D1 *Ebf1*+ and D2 *Stk32a*+) and the striosome (D1 *Ppm1e*+) is important because it suggests that voluntary cocaine taking engages both matrix- and striosome-associated MSN programs. Future efforts should seek to identify these-cocaine associated transcriptional adaptations in a non-activation dependent spatial context to help continue refining our understanding of the role of distinct MSN populations in cocaine related behaviors.

Astrocytes emerged as another major cocaine-responsive cell population with significant DEGs and cocaine-associated increase in IEG module score. These results align with astrocytes being increasingly recognized as active regulators of addiction-relevant synaptic plasticity rather than passive support cells. NAc astrocytes regulate extracellular glutamate, synaptic homeostasis, and drug seeking behavior [15, 16]. Additional work has shown that cocaine exposure induces functional adaptations in NAc astrocytes, that selective glia-mediated glutamate release can inhibit cue-induced cocaine seeking, and that astrocytic NMDA receptors in the NAc contribute to cocaine-induced conditioned place preference and associated neuroadaptations [38-40]. The present data add a cell type-resolved transcriptomic layer to this literature. Cocaine-associated astrocyte DEGs included *Hdac4, Pde10a, Pde7b, Cbs, Brwd1, Gse1*, and *Rab27b*, supporting chromatin regulation, cyclic nucleotide signaling, metabolic pathways, and vesicular or secretory mechanisms as candidate axes of astrocyte involvement in cocaine-induced plasticity. The predicted decrease in RAS/MAPK pathway activity in astrocytes, in contrast to the predicted increase in D1 *Ebf1*+ MSNs, further emphasizes that voluntary cocaine taking can engage the same broad signaling pathway, albeit in different directions across distinct cell types.

The partial mismatch between DEG results and IEG module-score results is another informative feature of the dataset. D1 *Ebf1*+ MSNs and astrocytes showed both prominent DEG burdens and cocaine-associated IEG module-score increases, suggesting that these populations were transcriptionally remodeled and in a heightened activity-associated state at the time of tissue collection. In contrast, oligodendrocytes showed a significant IEG module-score increase without being one of the top DEG-containing populations, whereas D2 *Stk32a*+ MSNs and D1 *Ppm1e*+ MSNs showed substantial DEG expression without comparably strong IEG module-score increases observed in D1 *Ebf1*+ MSNs. This dissociation suggests that some cocaine-associated MSN adaptations may reflect slower, non-canonical, or activity-independent transcriptional remodeling that would be missed by approaches focused only on immediate-early gene-defined neuronal ensembles. These patterns indicate that IEG induction and broader transcriptional remodeling capture are related but non-identical dimensions of cocaine response. This is consistent with recent ensemble-based studies showing that cocaine-context memories are transcriptionally encoded in Arc-defined NAc populations [41], while also demonstrating that important cocaine-associated adaptations occur outside classical IEG-defined ensembles. Thus, the current project should be useful not only for identifying activated populations, but also for discovering cell types undergoing molecular remodeling that may not be apparent from IEG expression alone.

This study provides an important and timely resource for the addiction field. The data identify D1 *Ebf1*+ MSNs as a dominant transcriptional substrate of voluntary cocaine taking. They also reveal substantial cocaine-associated remodeling in D2 *Stk32a*+ MSNs, D1 *Ppm1e*+ MSNs, and astrocytes, underscoring the value of resolving NAc biology beyond broad neuronal classes or regional averages. The convergence of synaptic, NMDA receptor-associated, RAS/MAPK, CREB-related, and astrocytic signaling programs provides a prioritized set of cellular and molecular mechanisms for future functional studies. More broadly, this project demonstrates how cell type-resolved transcriptomics can transform heterogeneous tissue-level cocaine-evoked responses into defined cellular hypotheses. By identifying the specific NAc cell populations and molecular programs engaged by volitional cocaine taking, these findings provide a foundation for mechanistic studies and, ultimately, the development of more precise therapeutic strategies for treating CUD.

## Acknowledgements

This work was supported by the following grants from the National Institutes of Health (NIH): R01 DA037897 and R01 DA061799 (HDS), R21 DA 057458 (HDS and RCC), and R21 DA 055846 (BCR); as well as institutional resources generously provided to BCR.

## Author Contributions

Study conceptualization: HDS, BCR.

Methodology: HDS, RM, BCR.

Software: RCC, JES, BCR.

Validation: HDS, RCC, SNC, BCR.

Formal Analysis: HDS, RCC, JES, BCR.

Investigation: HDS, RM, SNC, MF, VJ, JES, BCR.

Resources: HDS, BCR.

Data Curation: HDS, RCC, SNC, BCR.

Writing, Original Draft: HDS, RCC, BCR.

Writing, Reviewing and Editing: HDS, RCC, RM, SNC, MF, VJ, JES, BCR.

Visualization: HDS, RCC, BCR.

Supervision: HDS, BCR.

Project administration: HDS, BCR.

Funding Acquisition: HDS, RCC, BCR.

